# Beyond retinotopy: exploiting native visual representations in cortical neuroprostheses for vision loss remediation

**DOI:** 10.1101/2025.11.03.684808

**Authors:** Luca Baroni, Martin Picek, Saumil Patel, Andreas S. Tolias, Jan Antolik

**Affiliations:** Charles University, Faculty of Mathematics and Physics, Prague, Czech Republic; Department of Ophthalmology, Byers Eye Institute, Stanford University School of Medicine, Stanford, CA, US; Stanford Bio-X, Stanford University, Stanford, CA, US; Wu Tsai Neurosciences Institute, Stanford University, Stanford, CA, US; Department of Electrical Engineering, Stanford University, Stanford, CA, US

## Abstract

Cortical prosthetic systems offer a promising path to restoring vision in blindness by stimulating neurons in the visual cortex to evoke visual percepts. To effectively encode visual information, however, it is essential to target neurons based on their functional encoding properties. Current stimulation protocols focus on retinotopic information, ignoring other key encoding properties, and therefore fail to reproduce complex visual percepts. We demonstrate that incorporating orientation selectivity alongside retinotopy to guide stimulation dramatically improves fidelity of the evoked activity to the underlying neural code. We propose a Bottlenecked Rotation-Equivariant CNN (BRCNN) and demonstrate that neural responses can be predicted to a large degree using only retinotopy and orientation preference. Using this model, we design a retinotopy-and-orientation aware stimulation protocol and validate it in a state-of-the-art large-scale optogenetic stimulation simulation framework of primary visual cortex. Our protocol elicits neural activity patterns exhibiting substantially higher correlation with natural vision responses compared to retinotopic-only approaches.

## Introduction

Vision is our primary sense for navigating and interacting with the world, yet more than 40 million people worldwide suffer from blindness due to disorders and injuries affecting the visual system [8]. Visual neuro-prostheses (Fig. 1) aim to restore vision by bypassing non-functional parts of the visual system and directly write visual information directly into neural activity via neural stimulation [48,52]. These systems typically comprise three key components (Fig. 1B): a camera that captures visual input, a processing unit that converts input into stimulation patterns, and neurostimulators that delivers these patterns to target neurons (e.g., via electrodes or optogenetic actuators) [24,26]. To successfully evoke desired percepts, the processing unit must translate the visual input to stimulation patterns that drive neural activity, such that it is aligned with the intrinsic visual representations [7] (Fig. 1A-B).

**Figure 1.**
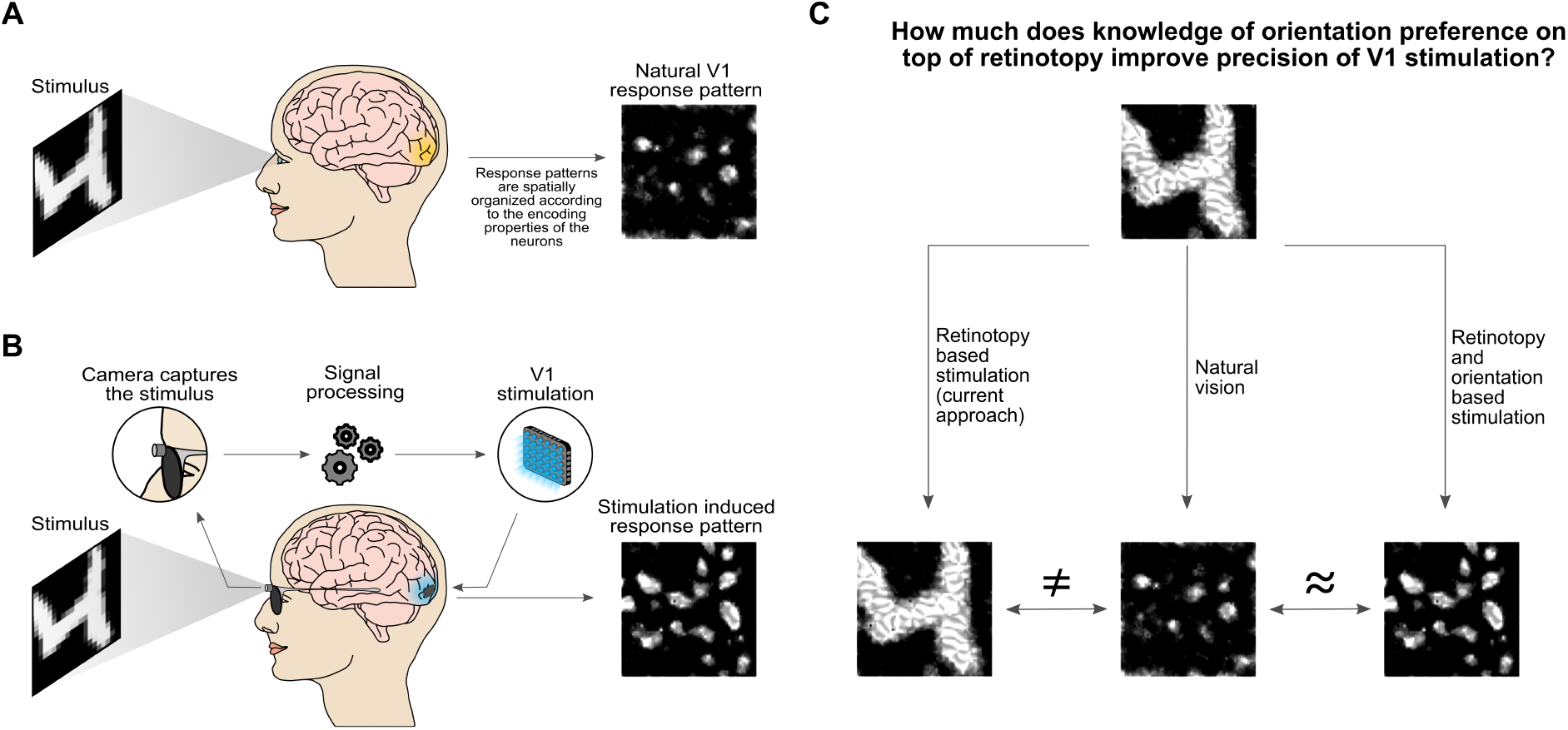
Conceptual overview of cortical prostheses and the role of neural encoding properties in V1 stimulation. (A) Natural vision: a visual stimulus elicits characteristic response patterns in V1, reflecting the encoding properties of neurons, with retinotopic position and orientation preference as the most prominent organizing principles. (B) Cortical prosthesis-based stimulation: a camera captures the stimulus, and a signal processing system generates stimulation patterns delivered to V1. To evoke meaningful percepts, an effective stimulation strategy should induce response patterns that resemble those occurring under natural vision conditions. (C) Importance of incorporating orientation preference: the potential improvement from including orientation preference in stimulation design. Left: a stimulation approach based solely on retinotopy produces a response pattern that deviates from natural vision. Right: incorporating both retinotopy and orientation preference leads to a response pattern that more closely matches natural vision.

While current prosthetic technologies cannot target individual neurons and instead activate local populations, the topographic organization of the visual system, where nearby neurons encode similar features [33,41], enables prosthetic stimulation to convey meaningful visual information by targeting local populations with shared coding properties. To date, visual prostheses have used this topographic organization to evoke phosphenes, localized discrete light percepts, and aimed to create simple composite visual percepts by activating multiple such phosphenes at appropriate coordinates in the visual field [4,18,25,47,60,63]. However, in current systems, such a phosphene-based approach limits the regained perception to the approximation of the target visual input by a limited set of dots (phosphenes). Visual prosthetics systems have been extensively studied in retina [25,47,63] for clinical cases where retina remains a viable target. Targeting of downstream areas, particularly the primary visual cortex (V1), has emerged as an attractive alternative due to its relatively low-level visual processing, and its favorable cortical magnification, allowing a denser coverage with stimulation elements [42] and possibility to address a broad range of visual impairments with one system. Crucially, beyond retinotopy, V1 exhibits additional exploitable topographically organized representations, most notably orientation preference [32]. Yet existing systems have entirely overlooked this organization. As a result, purely retinotopic stimulation strategies fail to capture key aspects of V1’s native code, and attempts to build complex percepts by summing phosphenes from multiple electrodes are bound to show significant limitations. Indeed, phosphenes induced by V1 stimulation have been reported to vary in their perceptual properties across electrodes, and simultaneous activation of multiple sites to often produce non-additive and inconsistent percepts rather than predictable composite images [27]. Incorporating additional encoding properties into stimulation protocols, particularly orientation preference, could thus yield activation patterns that better align with the brain’s natural visual representations (Fig. 1 C), potentially advancing visual neuroprosthetics beyond the ‘dotted vision’ paradigm, eliciting more naturalistic visual percepts.

This raises three key open questions we set out to address:

1. **How well can a retinotopic-and-orientation aware stimulation protocol recreate the visual code?** We first considered a theoretical best-case scenario in which neurons can be stimulated independently with arbitrary precision. We reformulated the question as a prediction task: we fit a 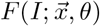 model that predicts responses to stimulus *I* for any V1 neuron using only its position 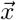 and preferred orientation *θ*, and take its predictive accuracy on V1 neurons from non-human primates as a conservative estimate of the best achievable fidelity under these assumptions.
2. **How can a retinotopic-and-orientation aware stimulation protocol be implemented?** Assuming a direct mapping between stimulation intensity and evoked neural activity, we use a trained 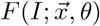 model to design a stimulation protocol where each element’s stimulation intensity is determined by the retinotopic position and orientation preference of the targeted neurons.
3. **How effective is such a protocol under realistic stimulation precision limits and network dynamics?** To address this question we used a previously published biologically detailed large-scale spiking model of V1 [2] enabling comparison of visually and optogenetically evoked responses while accounting for realistic stimulation precision and network dynamics. Using this model, we generated a synthetic dataset of stimulus-response pairs, fit an 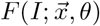 model to the data, designed a stimulation protocol from the fitted model, and evaluated it against a retinotopic stimulation baseline.

Addressing these questions yielded the following key findings:

a. A substantial portion of normalized response variability of V1 neurons from non-human-primates can be explained using only the knowledge of retinotopic position and orientation preference, despite neuron-to-neuron variations in other coding properties such as receptive field size, preferred spatial frequency, phase or non-linear spatial integration.
b. Our novel Bottlenecked Rotation-Equivariant Convolutional Neural Network (BRCNN) implementation of 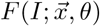 outperformed a simpler implementation based on the more established energy model [1] in predicting normalized V1 responses using only 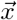 and *θ* as neuron-specific conditioning variables. This demonstrates that even in these constrained settings, data-driven models with higher representational capacity can capture encoding characteristics (such as contextual modulation) that more constrained classical parametric models miss.
c. The prediction accuracy of our BRCNN 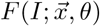 model generalizes to unseen neurons recorded from a different animal, indicating that it captures consistent species-wide functional structure of visual code. This demonstrates that our model trained on one subject can be used to inform stimulation strategies in others, facilitating clinical translatability of such stimulation strategies.
d. In our in-silico V1 stimulation experiments, our BRCNN stimulation protocol substantially outperforms a retinotopy-only approach corresponding to current stimulation strategies, producing activity patterns that much more closely resemble those evoked by natural vision. This demonstrates that the advantage of the retinotopic-and-orientation-based stimulation protocol persists even under the conditions of imperfect stimulation targeting inherent to current BMI systems. This highlights the importance of incorporating native encoding properties into stimulation strategies to generate meaningful response patterns and, consequently, more naturalistic percepts.

In summary, this study introduces a principled, data-driven approach for designing cortical stimulation strategies that, using encoding models, is firmly grounded in the native visual code. Providing 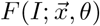, we make available a stimulation protocol that engages both retinotopic and orientation representations, and can be applied in preclinical animal studies and early human trials.

## Results

### Predicting macaque neural responses with BRCNN

In this section, we investigate how accurately neural responses can be predicted using only knowledge of neurons’ retinotopic positions and orientation preferences. To address this question, we developed a Bottlenecked Rotation-Equivariant Convolutional Neural Network (BRCNN) that naturally implements a function 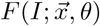, i.e. predicts neural responses based on these two properties alone.

The BRCNN consists of a rotation-equivariant convolutional core followed by a single-channel bottleneck (Fig. 2A). Rotation-Equivariant CNNs are a class of architectures introduced in prior work [19,23,64] extending the equivariance property of standard CNNs, originally limited to translations, to also include rotations. Whereas in CNNs the units in each channel extract features at different positions, in Rotation-Equivariant CNNs the units in each channel extract features at different positions and orientations. As such, their feature maps have four dimensions [*c, x, y, θ*] (corresponding to channels, positions, and orientations). By constraining the last layer of the core to be a single channel bottleneck (*c* = 1), we ensure that the same computation is applied at different positions and orientations across the feature map. Each neuron’s response is then predicted by reading from the position and orientation in this feature map corresponding to its retinotopic position and preferred orientation. Training the BRCNN to predict responses of a large population of V1 neurons, we implicitly learn the 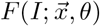 function that predicts neural responses using only position and orientation as varying parameters.

**Figure 2.**
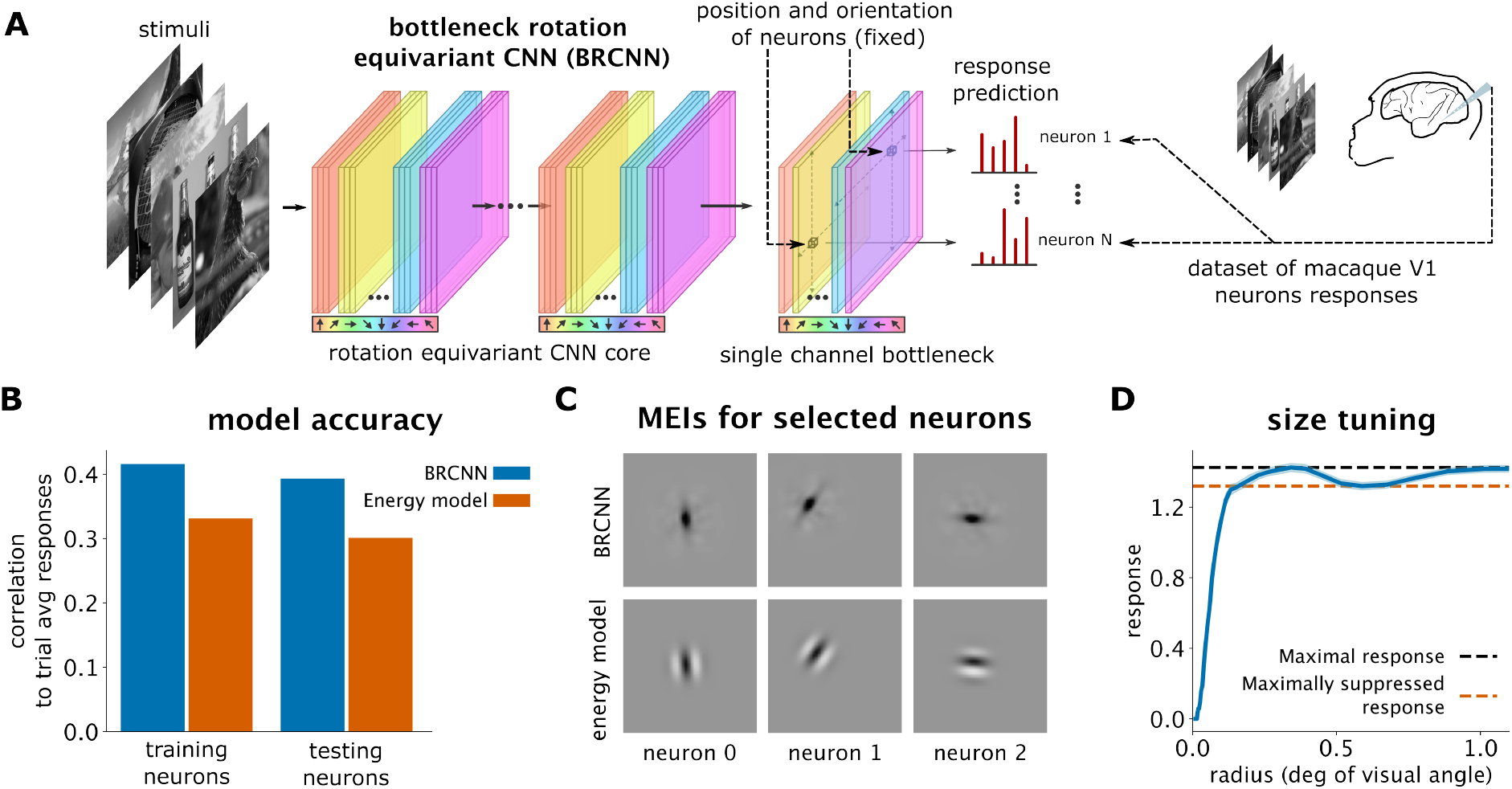
BRCNN predicts macaque neural responses: (A) Overview of the BRCNN model. The model takes static images as input and predicts neural responses. It features a rotation equivariant convolutional core whose units process information at different positions and orientations. The core’s final layer has a single channel, ensuring that each unit in the feature map performs the same computation at a specific position and orientation (different colors represent different orientations). A neuron’s response is predicted from the activation corresponding to the neuron retinotopic position and preferred orientation. Since all neurons read from the same single-channel feature map, they perform identical computations, differing only in position and orientation. (B) Bar plot showing the correlation between model predictions and trial-averaged neural responses to held-out images(blue: BRCNN; orange: energy model). Bars indicate mean correlation values for neurons used during training (left) and for a separate set recorded from a different macaque to assess generalization (right). (C) MEIs computed for three different neurons from BRCNN model and from energy model. (D) Size-tuning experiment response curve for neurons in BRCNN model.

As a benchmark, we fit the well-established energy model of V1 complex cells [1], which sums the squared outputs of a 90-degree phase-shifted pair of Gabor filters. Like the BRCNN, we restricted the model to depend only on retinotopic position and preferred orientation, with the remaining parameters shared across neurons.

We trained both models on a publicly available dataset of neurons recorded in the V1 of two macaques responding to greyscale natural images [12]. We used 238 neurons from one macaque for training and 214 neurons from the second macaque for evaluation, z-scoring each neuron’s responses (i.e., spike counts in a 120 ms windows 40 ms after stimulus onset) to account for firing rate differences. The BRCNN achieved significantly higher prediction accuracy than the energy model (Fig. 2B), outperforming it both on neurons from the first animal (evaluated on a held-out image set) and on neurons from the second. This ability to generalize to new neurons from a different animal with minimal performance degradation suggests that our model captures fundamental, species-wide properties of V1 neural encoding, rather than overfitting to specific neurons or subjects. Such generalization is key for eventual clinical applicability of our method. Notably, while our chosen architecture achieved strong generalization, we found that larger BRCNN variants tended to overfit to training neurons, highlighting the importance of adjusting model capacity to dataset size and diversity (see supplementary material).

To understand why BRCNN outperforms the energy model, we first computed Maximally Exciting Images (MEIs) for both models by optimizing input pixels under fixed contrast constraints [62]. As shown in Fig. 2C, BRCNN MEIs are identical across neurons except for position and orientation shifts, reflecting the single-channel bottleneck of our architecture. Notably, BRCNN MEIs differ from energy model predictions by preferentially assigning contrast to oriented OFF regions at the receptive field center, without prominent ON regions^1^, whereas the energy model does not have the flexibility to express such bias due to its construction. We hypothesize that this OFF preference reflects the known bias of V1 neurons toward darker stimuli [36,66].

We further hypothesized that the BRCNN’s superior performance can also be attributed to its ability to capture complex nonlinear encoding properties, which cannot be captured with an energy model, such as contextual modulation. To test this hypothesis, we conducted a virtual size-tuning experiment, presenting sinusoidal gratings of optimal orientation confined to circular aperture of increasing radius centered at each model neuron’s retinotopic position. The results (Fig. 2D) show that the BRCNN naturally learned both size-tuning and counter-suppression effects, matching known properties of V1 neurons [16]. The relatively low magnitude of the size-tuning likely reflects the fact that not all V1 neurons express size-tuning: since BCRNN cannot account for the inter-neuron variations of size-tuning magnitude, it predicts the averaged weak effect.

These results demonstrate that a substantial portion of V1 neural response can be explained through position and orientation information alone when implemented within an architecture capable of capturing complex nonlinear response properties.

### In-silico evaluation of BRCNN stimulation protocol for optogenetic BMI

Having demonstrated that neural responses can be effectively predicted using only retinotopic position and orientation preference, we next sought to evaluate such a stimulation protocol under more realistic conditions.

High-resolution visual prosthetic systems that will be capable of selectively targeting the orientation preference maps, such as those based on optogenetic [31] or infrared stimulation [59], are under active development. To determine, whether such future systems could effectively utilize our orientation-aware stimulation protocol, we performed an in-silico evaluation using a previously published simulation framework [2] that combines a large-scale biologically constrained spiking neural network model of primary visual cortex with a detailed model of optogenetic stimulation (Fig. 3B). A recent series of studies demonstrated that this model accurately replicates a broad range of visually and optogenetically evoked phenomena [2,3,13,53,58]. A key advantage of this simulation environment is that it enables direct comparison between visually and optogenetically evoked activity, while also accounting for light dispersion in tissue and recurrent network dynamics, which are critical limitations for the precision of targeted optogenetic stimulation in V1. Together, these features make it an ideal testbed for evaluating the effectiveness of stimulation protocols against ground-truth responses that are not yet experimentally obtainable in columnar species during stimulation at single-neuron resolution at the multi-millimeter spatial scale of cortical maps. In this environment, we quantitatively assessed the performance of a retinotopy-andorientation-aware BRCNN stimulation protocol against a retinotopic-only protocol representing current stimulation strategies.

**Figure 3.**
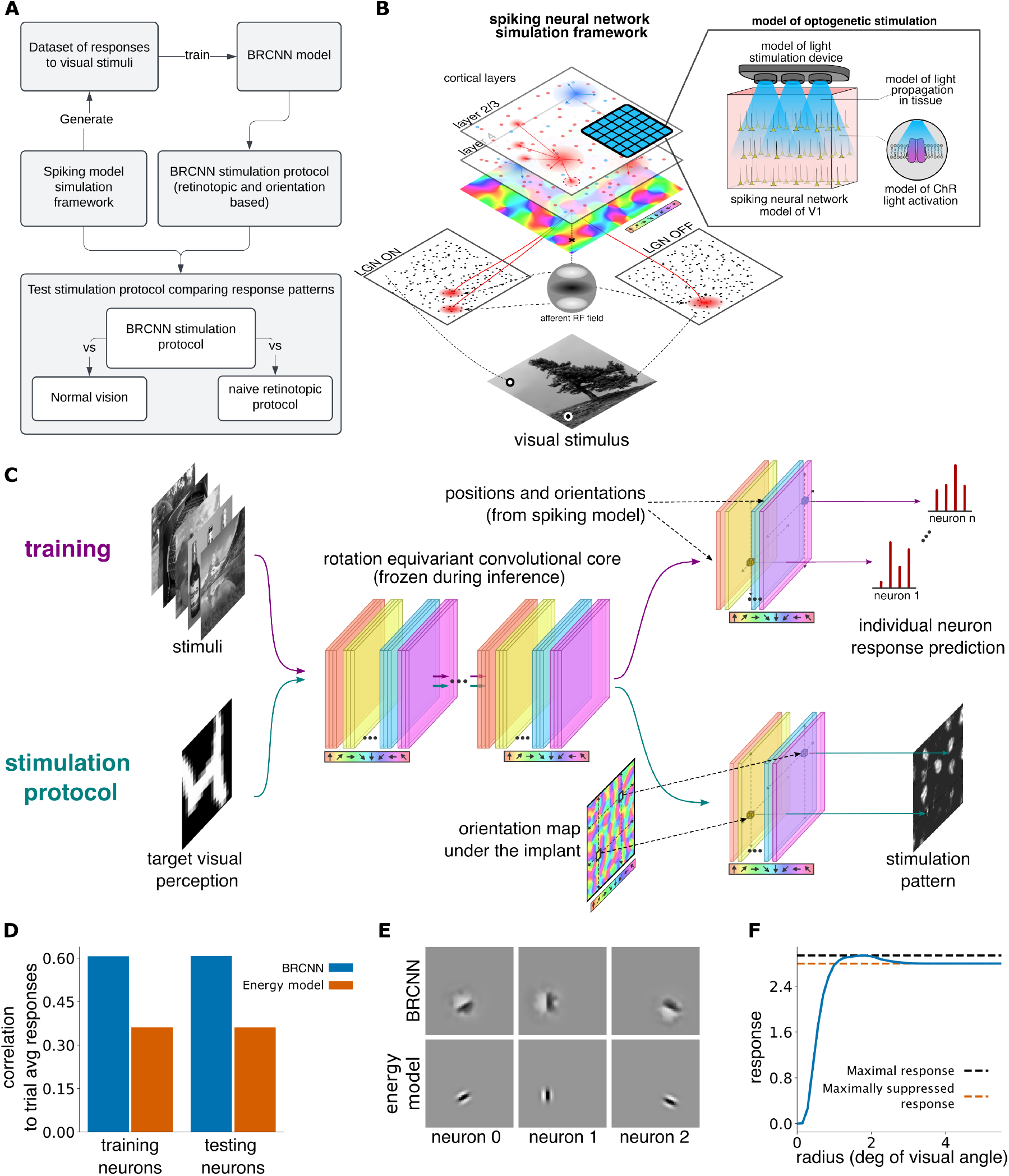
Computational framework for designing and validating the BRCNN protocol. (A) Diagram illustrating the pipeline used for designing and evaluating the BRCNN stimulation protocol. A large-scale spiking model generates a dataset of neural responses to set of visual stimuli, which is then used to train the BRCNN model. The trained BRCNN model is subsequently used to implement a optogenetic stimulation protocol, which is tested within the same simulation framework. The quality of the stimulation protocol is then assessed by comparing the optogenetically vs visually evoked response patterns to the same target visual stimulus. (B) Overview of the simulation framework. It consists of a large-scale spiking neural network model with center-surround LGN ON and OFF layers and two cortical layers containing excitatory and inhibitory neurons: Layer 4, whose excitatory neurons receive input from the LGN, and Layer 2/3, whose excitatory neurons can be optogenetically stimulated. Optogenetic stimulation is modeled through a simulation of the light stimulation device, light propagation in tissue, and ChR-mediated neural activation. This simulation framework allows comparison of responses across different stimulation conditions. (C) The BRCNN models are first trained and then used as a stimulation protocol, providing the target optogenetic activation levels given the knowledge of the retinotopic position and orientation preference of the neurons beneath the implant. (D) Bar plot showing the correlation between model predictions and trial-averaged neural responses to held-out images (blue: BRCNN; orange: energy model). Bars indicate correlation values for neurons used during training (left) and for a held-out set (right). (E) MEIs computed for three different neurons from BRCNN model and from energy model. (F) Size-tuning experiment response curve for neurons in BRCNN model.

Fig. 3A outlines the methodology we followed in this section. First, we generated a large dataset of neural responses to visual stimuli from all neurons in the spiking network model. Then, following our approach on macaque data, we trained a BRCNN and an energy model on responses from a subset of these neurons and evaluated generalization performance on the remaining held-out neurons. The BRCNN trained on simulated responses achieved a correlation of 0.61 on both training and held-out neurons, significantly outperforming the energy model (Fig. 3D). This higher correlation and perfect generalization observed in the simulation, compared to macaque data, likely stems from the lower intrinsic neural variability in the spiking model and the larger dataset size (25,000 vs. 238 neurons; 100,000 vs. *≈* 24,000 training images ^2^). Additional MEI and size-tuning analyses confirmed that the BRCNN captured oriented features and surround suppression (Fig. 3E-F).

We used the BRCNN’s 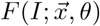 mapping to implement a stimulation protocol where each stimulation element activates according to the retinotopic position and orientation preference of its underlying neural tissue (Fig. 3C). We defined the stimulation strength 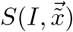 at cortical position 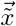 for input image *I* as:

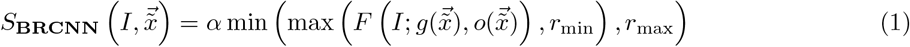

where 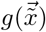 and 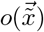 correspond to the mapping of retinotopic position and preferred orientation onto position along cortical surface, respectively. The parameter *r*_*min*_ sets a threshold to avoid stimulation in weak activation regions, while *r*_*max*_ caps the maximum stimulation intensity. The factor *α* is an arbitrary linear scaling parameter converting predicted response magnitudes to stimulation intensity (see Methods for details). To assess the advantage of this orientation-aware BRCNN-based protocol, we compared it to a naive retinotopic baseline in which stimulation strength at a given cortical location 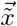 is directly proportional to local luminosity at the corresponding retinotopic position 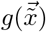:

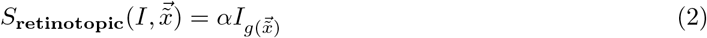

We tested both protocols using two types of visual stimuli: oriented gratings to facilitate interpretability and MNIST digits to evaluate human-centric percepts [40]. For oriented gratings (Fig. 4A), responses under natural vision conditions show selective activation of neurons based on their preferred orientation. Our BRCNN protocol successfully reproduced this selective activation pattern, while the retinotopic protocol activated neurons indiscriminately across all orientation preferences. For MNIST digits (Fig. 4B), natural vision responses engaged different orientation domains according to the location of oriented features in the stimulus. Even in this case, our BRCNN protocol generated response patterns closely reflecting those of natural vision, whereas the retinotopic protocol continued to produce non-selective activation patterns that poorly matched the target responses. To quantitatively evaluate these results, we computed, for each stimulus, the Pearson correlation coefficient between natural vision and optogenetically evoked responses across all neurons in the network. Our BRCNN protocol generated activity patterns that strongly correlated (*r ≈* 0.5) with natural vision responses, maintaining comparable performances across both oriented gratings and more complex MNIST stimuli (Fig. 4C). In contrast, the retinotopic protocol produced activation patterns that showed negligible correlation with the tar get responses.

**Figure 4.**
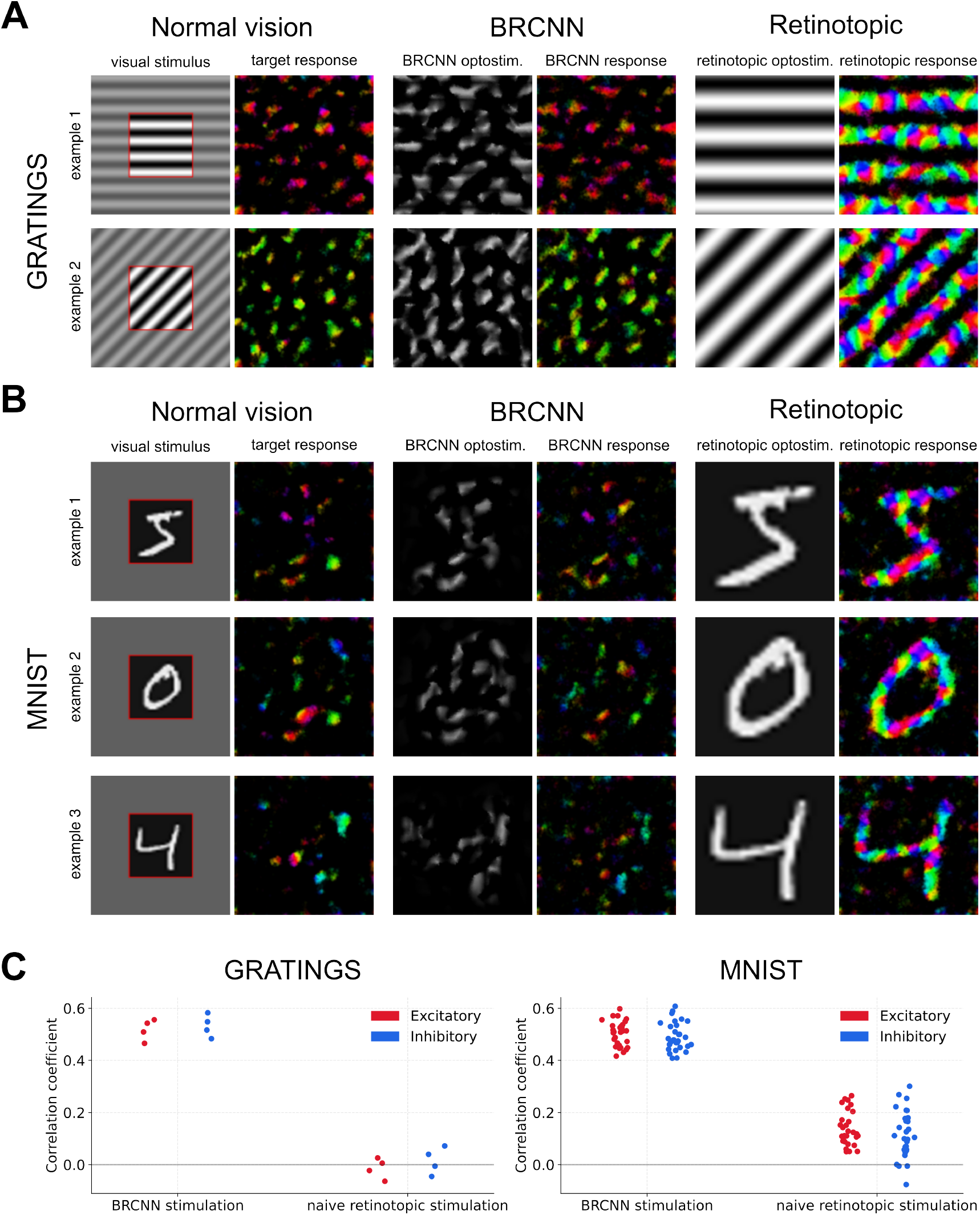
Comparing BRCNN-based and retinotopy-based stimulation protocols. **A, B**, Comparison of response patterns for different stimulation protocols. Each row corresponds to an example stimulus presented to the spiking neural network model either under normal vision conditions or using optogenetic stimulation protocols. The two leftmost columns show: (1) the visual stimulus and (2) the corresponding visually evoked neural response, averaged across 20 trials. The red square in the stimulus indicates the region of the visual field covered by the neurons’ retinotopic positions in the model, with stimulus contrast outside this region reduced for illustrative purposes. The two center columns show the BRCNN-based stimulation protocol: (1) the optogenetic stimulation pattern generated by the BRCNN model and (2) the resulting optogenetically evoked neural activation. The two rightmost columns show the naive retinotopy-based stimulation protocol: (1) the optogenetic stimulation pattern generated by the retinotopic protocol and (2) the corresponding optogenetically evoked neural activation. **A** Grating stimuli. **B** MNIST stimuli. **C** Quantitative evaluation of stimulation accuracy. Scatter plots show the correlation coefficients between model-predicted and target neural responses for excitatory (red) and inhibitory (blue) neurons. The BRCNN-based stimulation outperforms naive retinotopic stimulation across both grating and MNIST stimuli. In these scatter plots, each neuron is represented by a dot whose color indicates its preferred orientation and whose transparency reflects its firing rate (bright color = higher rate)

Overall, our in-silico experiments show that a retinotopy-and-orientation-aware BRCNN stimulation protocol recovers natural vision-like, orientation-selective activity in V1 under realistic optical and network constraints, with substantially higher correlation to natural-vision responses than the retinotopic baseline. These findings demonstrate the feasibility of orientation-informed stimulation and underscore its importance in the design of stimulation protocols for cortical prostheses.

## Discussion

In this study we demonstrate, for the first time, how incorporating knowledge of two fundamental organizing principles of V1, retinotopy and orientation selectivity, can dramatically improve the fidelity of cortical prosthetic stimulation for vision restoration.

Using a BRCNN that implements a stimulus-to-response function 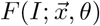, we demonstrate that neural responses can be surprisingly well predicted using only information about the position and orientation of their receptive field. The BRCNN outperforms simpler alternatives, such as energy models, due to its capacity to learn complex V1 response properties including surround suppression. Importantly, the model’s performance generalizes across different subjects. This makes it particularly suitable for prosthetic applications where protocols can be trained on high-quality data from initial cohorts of patients (or possibly even from primate models, which share very similar visual coding, and where substantially higher data quality and volumes are achievable) and then deployed across a broader patient population without requiring individual data collection and training procedures. Then, we tested the effectiveness of a retinotopy-and-orientation-aware BRCNN-based stimulation protocol within a simulation framework that allows controlled comparison of different optogenetic stimulation strategies while accounting for factors that limit optogenetic BMI precision, such as light dispersion and secondary recurrent network activation. Our results clearly demonstrate that it substantially outperforms standard retinotopic-only stimulation methods, yielding cortical activity patterns that much more faithfully replicate those seen under natural vision.

While our findings indicate a clear path toward improving the effectiveness of stimulation protocols by incorporating orientation selectivity, several important challenges and considerations must be taken into account when interpreting our results. A key issue for clinical translation is that functional maps, such as retinotopy and orientation preference, are typically derived from visually evoked responses, which are not available in blind patients. Since our method relies on these maps, alternative reconstruction techniques are needed. Importantly, retinotopic position has been reconstructed from spontaneous activity in animals and humans [44], and animal studies indicate that orientation preference maps can likewise be inferred [37,56]. While the accuracy of orientation-map reconstruction in humans remains to be established, particularly in long-term blindness, further research could refine these approaches and enable reliable patient-specific mapping for visual prosthetics [39]. Additionally, while our results demonstrate generalization across macaques, inter-species differences in V1 organization might still require adaptation of the BCRNN model for human cortical stimulation. Regardless, given the shared V1 coding properties in primates, models trained on non-human primate data hold strong potential for guiding effective human cortical stimulation. Another translational challenge relates to stimulation modality. Current human cortical implants rely on electrical stimulation which has very low spatial precision due to the predominant activation of axonal processes [29]. Our study evaluated stimulation using an optogenetic stimulation framework due to its potential to stimulate the cortical surface with enough spatial precision to selectively target the topologically organized orientation preference maps in V1. However, translation of such advanced optogenetics-based BMIs into clinical applications is still under development.^3^.

Regardless of stimulation modality, its effective resolution, determined by density and size of the stimulation elements, propagation properties of the stimulation signal, and neural network effects, is critical for taking advantage of the intrinsic cortical organization. Encouragingly, recent advances in prosthetic technologies are steadily improving stimulation precision, paving the way for more effective prosthetic vision [31,59]. Yet, high-resolution stimulation alone is insufficient; it must be coupled with stimulation protocols that effectively harness the underlying visual representations. To this end, we provide the first ready-to-use visual representation-aware stimulation protocol that can be tested in the new generation of high-precision BMI systems.

Finally, while our approach focuses on retinotopy and orientation selectivity, V1 neurons also encode additional topographically organized properties, such as color selectivity and spatial frequency tuning [15,34,43]. In this initial study, we prioritized retinotopy and orientation selectivity as they remain the dominant properties of V1 representations and are currently the most feasible to extract from spontaneous activity. Nevertheless, targeting neurons based on additional topographic properties could further improve stimulation fidelity, showing future directions for developing our stimulation approach.

Acknowledging these challenges, we suggest that future work should advance along two complementary paths. On the technical side, our BRCNN could be improved by enlarging the training set and restricting training data to neurons from layers targeted by stimulation (e.g., superficial layers for surface optogenetics). Additionally, our approach could be extended to process video inputs, enabling dynamic stimulation patterns that better capture the temporal aspects of natural vision. In parallel, experimental validation should progress towards clinical applications. Behavioral experiments in animal models, such as shape discrimination tasks [18], could assess whether orientation-aware stimulation enables better artificial visual percepts than retinotopic-only approaches. Moreover, once methods for reconstructing orientation maps from spontaneous activity in blind humans are sufficiently developed [39], our protocol should advance to preclinical human trials. These studies could directly assess whether patients report more consistent, interpretable visual percepts when orientation selectivity is incorporated into stimulation protocols.

## Methods

### Bottlenecked Rotation-Equivariant CNN (BRCNN)

The Bottlenecked Rotation-Equivariant CNN (BRCNN) architecture consists of two main components: a rotation-equivariant convolutional core [19,23,64] for feature extraction and a specialized readout layer for neural response prediction. Visual stimuli are processed through the equivariant core to produce rotation-equivariant feature representations, which are then used by the readout to predict the responses of recorded neurons based on their retinotopic position and preferred orientation. In the following paragraphs, we describe in detail the components of the BRCNN model, while providing a brief introduction to the rotation-equivariant convolutions used in its core. For a more mathematical and comprehensive discussion see Cohen and Welling [19].

**BRCNN core** Rotation equivariant convolutions are an extension of standard convolutions defined to extend their property of equivariance to translations to include both rotations and translations (formally speaking, forming the SE(2) group).

Standard convolutions are said to be equivariant to translations as they satisfy the following property:

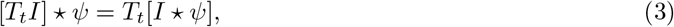

where *I* is an input, *ψ* is the convolution filter (or kernel), and *T*_*t*_ is the translation operator acting as 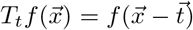. When implemented as neural network layers, standard convolutions outputs feature maps indexed by channel and discrete position indices (*x* and *y* for 2D inputs and convolutions). Each channel corresponds to a specific computation and the position indices (*x, y*) indicate where in the input this computation is applied.

Similarly, when rotation equivariant convolutions are implemented as neural network layers, they output feature maps with channel, discrete position *x, y*, and discrete orientation indices *θ*. For each fixed channel and orientation index, the same computational operation is applied at each spatial location (*x, y*). When varying the orientation index *θ*, the same computation is applied but rotated accordingly.

Rotational equivariant CNNs are implemented with two types of layers. The first layer is a lifting layer that maps input images into structured feature maps with channel, height, width, and orientation indices. Then, rotational equivariant convolutional layers further process these feature maps, preserving their structure.

The lifting layer performs the following operation

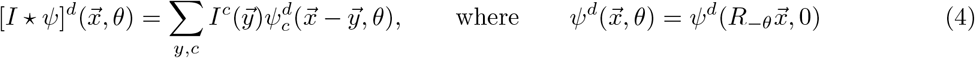

For each output channel *d*, the input *I* is convolved with rotated versions of the same base filter 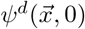 to generate the various orientations *θ* in the output feature map. This constraint is necessary to preserve equivariance to rotations.

The subsequent rotation equivariant convolutional layers process these orientation-structured feature maps performing the following operation:

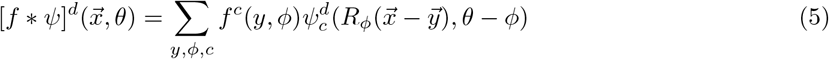

which implies that (a) for any output channel *d* and output orientation *θ*, as many spatial filters as orientations and input channels are convolved with the corresponding spatial feature maps and then summed together; (b) when varying the output orientation, the same filters are spatially rotated and cyclically permuted before being applied to the same feature maps.

Following the above outlined principles, the core of the BRCNN model is implemented with an initial lifting layer followed by standard rotationally equivariant convolutional layers. We modify standard rotation-equivariant CNNs by introducing a single-channel bottleneck (*c* = 1) in the final layer, enforcing shared computations across the feature map, differing only in the positions and orientations at which they are applied.

To represent filters at multiple spatial orientations (other than 0, 90, 180, and 270 degrees) while avoiding aliasing artifacts, we represented them in a steerable basis of functions [64]. Rather than learning the pixel values of filters directly, this approach learns the coefficients of a linear combination of 2D basis functions to construct each filter. Following Ecker et al. [23], we use 2D Hermite functions in polar coordinates [30,61], employing functions up to rank *k* for filters of size *k*.

#### BRCNN readout

The readout layer predicts neurons’ responses from specific positions and orientations in the core’s output feature map, based on pre-estimated retinotopic positions and preferred orientations of the modeled neurons. Values between discrete positions and orientations are predicted linear interpolating nearby values. We applied periodic boundary conditions to possible orientations values.

### Energy model

The energy model is a well-established model of V1 complex cells, which are the predominant cell type in Layer 2/3. It is implemented by summing the squared outputs of a 90-degrees phase-shifted pair of Gabor filters:

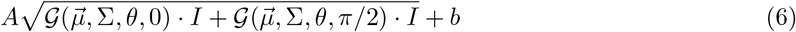

where 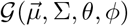 is a Gabor filter with parameters 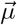 (the center of its Gaussian envelope), Σ (determining the spread of its Gaussian envelope), *θ* (determining its orientation), and ϕ (determining its phase, set to 0 and *π/*2 in the energy model). In our implementation, we constrained the parameters 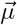 and *θ* to correspond to the retinotopic position and preferred orientation of each recorded neuron, while learning all remaining parameters as shared parameters across the population. In all experiments involving an energy model the input images were linearly rescaled so that the mid-scale gray value was set to 0.

### Optogenetic V1 Model Simulation Framework

#### Spiking Neural Network Architecture

The large-scale V1 model is described in detail in [3]. It is a model of two 5 *×* 5 mm patches of cortical layers, Layer 4 and Layer 2/3 specifically, based on experimental data from cat V1. At an eccentricity of roughly 3 degrees, cortical magnification factors is 1 and therefore neurons receptive fields covers roughly 5 *×* 5 degrees of visual field. Each cortical layer contains 46875 cortical neurons, divided into two populations, excitatory and inhibitory, with a ratio of 4:1, consistent with biological findings.

The Layer 4 populations receive thalamic input from two populations of neurons (ON-centered and OFF-centered) that model retinal/LGN input. The model captures both the feed-forward pathway (where external visual stimuli are processed by retinal/LGN neurons before reaching V1) and the recurrent cortical V1 pathway. Intra-cortical connectivity dominates the number of connections, with only 14% coming from thalamic input, consistent with anatomical findings. In line with anatomical findings, all cortical populations present short-range connectivity, while Layer 2/3 excitatory neurons additionally present long-range connections to other L2/3 excitatory and inhibitory neurons, and excitatory Layer 4 neurons send narrow projections to both excitatory and inhibitory Layer 2/3 neurons.

#### Cortical neuron model

All cortical neurons are modeled as single-compartment exponential integrate-and-fire units.

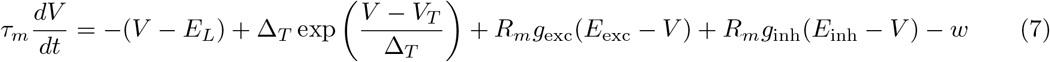

When the membrane potential reaches a threshold of *−*40 mV, an action potential is triggered and the membrane potential is reset to *V*_*r*_ = *−*60 mV during a refractory period set to 2*ms* for excitatory neurons and 0.5 ms for inhibitory neurons. The membrane time constant *τ*_*m*_ is set to 8*ms* for excitatory cells and 9 ms for inhibitory cells. The leak potential *E*_*L*_ and firing threshold *V*_*T*_ are set to *−*80 mV and *−*57 mV for excitatory neurons, and *−*78 mV and *−*58 mV for inhibitory neurons. The slope factor Δ*T* is set to 0.8 mV across all neurons. Membrane resistance *R*_*m*_ is set to 250 MΩ for excitatory neurons and 300*M*Ω for inhibitory neurons. Excitatory and inhibitory reversal potentials *E*_*exc*_ and *E*_*inh*_ are set to 0 mV and *−*80 mV respectively.

#### Retina/LGN input model

Retinal/LGN neurons (from here refer to as simply LGN neurons) are modelled as conductance based leaky integrate-and-fire. Their response is driven by injected currents obtained by first convolving spatiotemporal visual stimulus with the spatiotemporal receptive field of the given LGN neuron constructed as a difference-of-Gaussians (ON or OFF centered) spatial profile multiplied by a bi-phasic temporal profile defined by a difference of gamma functions. Exact spatial and temporal parameters were fitted to match the experimental observations in Cai et al. [14]. Subsequently, the output of the convolution is scaled by contrast and luminance to match the experimental observations of contrast and luminance saturation reported in Bonin et al. [6], Papaioannou and White [50]. Please refer to Antolík et al. [3] for exact detailed of the fitted model parameters. The resulting traces are then injected into the LGN neurons, together with an additional white noise current whose magnitude and variance are set such that both ON and OFF LGN neurons fire at the experimentally recorded rates under spontaneous conditions, respectively *≈* 17 and 8 spikes/s according to Papaioannou and White [50].

As the cortical layers span 5 *×* 5 degrees of visual field, to accommodate the full extent of the receptive field of neurons at the edges of the model, the LGN population covers a larger area measuring 6 *×* 6 degrees of visual field. Similarly, to accommodate the whole spatial extent of the spatiotemporal receptive field of the LGN neurons, the simulated visual field spans 11 *×* 11 degrees.

#### Thalamo-cortical connections

Each Layer 4 excitatory and inhibitory neuron receives input from both ON- and OFF-centered LGN populations [20]. For each cortical excitatory neuron the number of incoming ON and OFF connections is randomly drawn from the range [45, 95], whereas for inhibitory neurons the range is [56, 84]. For each Layer 4 neuron, the connections from the ON and OFF neurons are then probabilistically sampled according to a spatial connectivity distribution modeled as the sum of a half-wave rectified Gabor function and a two-dimensional Gaussian superimposed on the ON and OFF sheets.

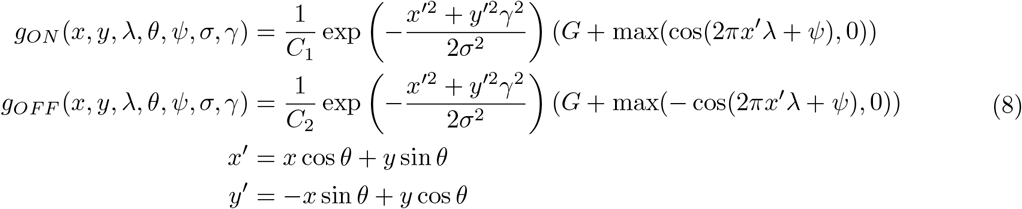

with *G* = 0.085 representing the relative weight of the Gaussian with respect to the Gabor distributions and *C*_1_, *C*_2_ are normalization constants. The neuron’s phase preference *ψ* is random. To induce functional organization in the model, the orientation *θ* of the Gabor distribution is obtained from a pre-computed 5 *×* 5 degrees orientation map overlaid onto the model cortical surface. For simplicity, the remaining parameters are set to constant values. In line with experimental findings, the spatial frequency *γ* was set to 0.8 cycles per degree [46], the aspect ratio of the Gabor distribution *γ* to 2.5 [51] and the size *σ* to 0.17 [65].

#### Cortico-cortico connectivity

Cortico-cortical connectivity is structured as follows: each Layer 4 excitatory neuron receives 1000 cortico-cortical synapses, while Layer 2/3 excitatory neurons receive 2300 synapses. Inhibitory neurons in both layers receive 40% of the synapses compared to their excitatory counterparts, accounting for their smaller size. Synapses are sampled according to the population density ratio of 4:1 (excitatory:inhibitory). Of the synapses targeting Layer 2/3 neurons, 22% of the excitatory connections originate from Layer 4 excitatory neurons. Additionally, the model includes a direct feedback pathway from Layer 2/3 to Layer 4, which comprises 20% of the Layer 4 synapses. While such direct connections are rare, strong feedback from Layer 2/3 to Layer 4 via layers 5 and 6 exists [5], and this loop appears to be necessary for correct functional properties across layers [3].

The spatial organization of the cortico-cortical connections follows two principles: 1) the connection probability decreases with cortical distance [10,11,57], and 2) connections present specific functional biases [11,38]. To combine these two principles into a single connection probability density function, separate probability functions are computed for each principle, which are then multiplied and renormalized.

For all intra-layer connections, with the exception of connections emanating from Layer 2/3 excitatory neurons, and feed-back excitatory connections from *L*2*/*3 *→ L*4, the spatial component follows a hyperbolic distribution

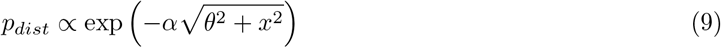

where *x* represents the distance between two neurons, *α* and *θ* are parameters obtained by fitting connection probability data in Stepanyants et al. [57] (see Antolík et al. [3] for a comprehensive report of the parameters). The intra-cortical functional specificity of Layer 4 connections is organized according to a push-pull scheme, with excitatory neurons being more likely to form connections with neurons with correlated thalamo-cortical receptive fields (Eq. 8) and inhibitory neurons more likely to connect with neurons with anticorrelated thalamo-cortical receptive fields.

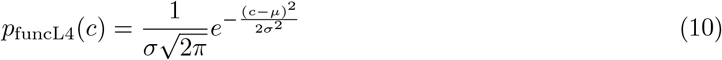

where *c* is the correlation between the two neurons thalamo-cortical receptive fields, *σ* = 1.3 and *µ* = *±* 1 depending on whether the pre-synaptic neuron is excitatory or inhibitory. Feedforward (*L*4 *→ L*2*/*3) excitatory connections functional component is given by

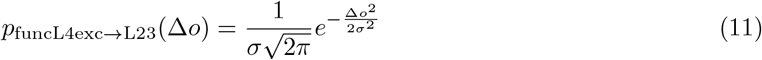

biasing connections to be formed between neurons with similar orientation preference. Here Δ*o* is the difference of orientation preference between the pre-synaptic and post-synaptic cells based on the precomputed orientation map and the standard deviation is set to *σ* = 1.3.

According to [11], in Layer 2/3, excitatory neurons form both local connections and long-range functionally biased connections. Accordingly, we modeled the connection probability as:

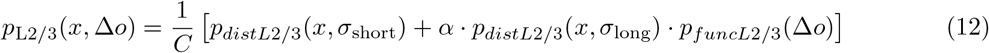

where *p*_*distL*2*/*3_(*d, σ*) are Gaussian distributions over distance:

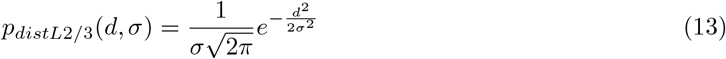

and *p*_*funcL*2*/*3_(Δ*o*) is a Gaussian function of orientation preference difference:

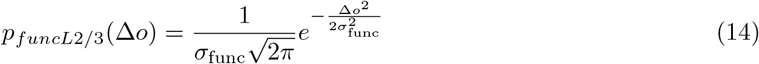

Here, *C* is a normalization constant, *α* represents the relative strength of long-range connections, *x* is the distance between neurons, and Δ*o* is the difference in orientation preference between pre- and postsynaptic cells. The parameters were set to *σ*_short_ = 270 *µ*m, *σ*_long_ = 1000 *µ*m, *σ*_func_ = 1.3 rad, and *α* = 4, based on fits of the model to cat data [11]. Conversely, Layer 2/3 inhibitory neurons connections were modeled with no functional specificity (*p*_*funcL*2*/*3*inh*_ = 1).

Finally, feedback connections (*L*2*/*3 *→ L*4) are modeled as local and functionally specific and obtained as the normalized product of the distributions in equations 13 and 14 with *σ*_*dist*_ set to 100 *µ*m and *σ*_*func*_ set to 1.3 radians for excitatory post-synaptic neurons and 3 radians for inhibitory post-synaptic neurons.

#### Synapses and delays

All Synapses in the model are implemented as exponential conductance changes

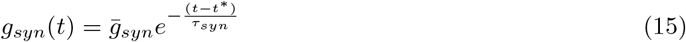

where *t*^*\**^ is the spike time,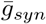 represents the synaptic weight, and *τ*_*syn*_ is the decay time constant set to 1.5 ms for excitatory synapses and 4.2 ms for inhibitory synapses. Synaptic weights 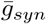 vary by connection type: Layer 4 excitatory-to-excitatory (0.18 nS), Layer 4 excitatory-to-inhibitory (0.22 nS), inhibitory synapses (1 nS), Layer 2/3 excitatory-to-inhibitory (0.35 nS), thalamo-cortical (1.2 nS), and feedforward Layer 4 to Layer 2/3 (1 nS).

Short-term synaptic depression is implemented according to the Tsodyks-Markram [45] model, with distinct parameter sets assigned to different connection types. All synapses use a utilization factor (*U* = 0.75) and no facilitation (*τ*_fac_ = 0 ms). Excitatory synapses have a postsynaptic decay time constant (*τ*_psc_ = 1.5) ms. Thalamo-cortical synapses are modeled with strong depression (*τ*_rec_ = 125 ms), intra-layer and feedforward (*L*4 *→ L*2*/*3) excitatory synapses with moderate depression (*τ*_rec_ = 30 ms), and feedback connections (*L*2*/*3 *→ L*4) with weaker depression (*τ*_rec_ = 20 ms). Inhibitory synapses have longer decay constants (*τ*_psc_ = 4.2 ms) and layer-specific recovery times: 70 ms in Layer 4 and 30 ms in Layer 2/3.

Synaptic delays combine a distance-dependent propagation component set to 0.3 ms^*−*1^ for intra-cortical connections [9, 17] with an additive, connection-type-specific constant delay: 1.4 ms for excitatory to excitatory, 0.5 ms for excitatory to inhibitory, 1.0 ms for inhibitory to excitatory, and 1.4 ms for inhibitory to inhibitory connections. This additive component reflects experimentally observed differences in synaptic transmission latencies between neuron types [49]. Thalamo-cortical delays are independently sampled from a uniform distribution between 1.4 and 2.4 ms.

#### Light emitting elements and light propagation in cortical tissue models

Following Antolik et al. [2], we simulate optogenetic stimulation using a regular 5 *×* 5 mm lattice (pitch 10 *µ*m) of identical circular light sources placed on the cortical surface, targeting Layer 2/3. The maximum light intensity at the cortical surface across the array is set to 3.0 *×* 10^14^ photons*/*s*/*cm^2^.

To simulate the effect of light attenuation with depth, we assigned each neuron a depth coordinate *d*_*n*_, in addition to its lateral position 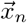. This depth is used exclusively for computing light flux; all other aspects of the simulation (e.g., connectivity and delays) depend only on lateral positions. Light propagation in cortical tissue was simulated using the ‘Human Brain Grey Matter’ model in LightTools, assuming a 590 nm emission wavelength. Scattering and absorption are modeled using the Henyey-Greenstein phase function with anisotropy factor *g* = 0.87 and mean free path 0.07 mm, based on Jacques [35]. These simulations generated a 2D lookup table *T* (*d, l*) that captures the relative light flux (photons/s/cm^2^) at a given tissue depth *d* and lateral distance *l* from the source. Using this lookup table, the total light flux *γ*_*n*_ at the location of neuron *n* is computed as a linear sum of the contributions from each emitting element *e*:

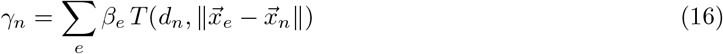

where *β*_*e*_ is the light flux of emitter *e* at the cortical surface, and 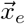 and 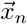 are the lateral positions of emitter *e* and neuron *n*, respectively.

#### Channelrhodopsin model

We follow the same modeling approach for ChrimsonR channel dynamics as in Antolik et al. [2], using the Markov kinetic model introduced by Sabatier et al. [54]. The model represents the protein as a five-state Markov system [22], where each state corresponds to a different conformation of the ChrimsonR channel. Transitions between states can be thermal (with fixed time constants) or photo-induced (with time constants depending on the current light intensity and occurring only in the presence of light). The evolution of state occupancies is governed by a linear system of differential equations depending on the transition time constants and the light stimulus. The resulting membrane conductance is computed from the number of channels in the open states and their respective conductances. Parameter values were taken from Sabatier et al. [54], who developed and fit the model to patch-clamp recordings from ChrimsonR-expressing HEK293T cells under 590 nm light stimulation.

### Datasets, Neuron Parameter Estimation and Model Trainings

#### Macaque V1 Dataset

The macaque experiments in this work rely on a dataset made publicly available by **?**]. It contains extracellular recordings from 458 V1 neurons collected across 32 sessions (17 from one animal and 15 from another), using a 32-channel linear electrode array spanning multiple cortical layers. Recordings were obtained from awake, fixating rhesus macaques. The stimulus set consists of grayscale natural images sampled uniformly from the ImageNet dataset, each spanning 6.7° of visual angle. Images were presented for 120 ms without interleaving blanks and were centered at the retinotopic location of the recorded neurons. Neuron eccentricities ranged from 2° to 3°. The dataset includes a training set consisting of 24000 natural images presented once across multiple sessions, and a test set comprising a separate subset of 75 images shown in each session, each repeated for 40–50 trials. Neuronal responses were extracted by counting the number of spikes, after spike sorting, within a time window from 40 ms to 160 ms following stimulus onset.

#### Macaque V1 Neurons Retinopic positions and Preferred Orientation Estimations

To estimate the retinotopic position and preferred orientation of the macaque neurons, we used the V1 model from Fu et al. [28] fitted to the macaque data used in this study. Following their approach, we performed two in silico experiments on the model: a sparse noise stimulation protocol to estimate receptive field (RF) location, and an orientation tuning experiment to determine preferred orientation. The model takes as input images of 93×93 pixels, corresponding to 2.67° of visual angle. Stimuli for the sparse noise experiment consisted of white or black squares of 4×4 pixels (corresponding to 0.11×0.11°) presented on a mid-scale gray background. To remove polarity and isolate excitatory regions, we first constructed a polarity-agnostic version of the stimuli by mapping black squares to white squares, effectively treating black and white stimuli as equivalent. We then averaged the corresponding responses after subtracting the baseline response (measured during gray background stimulation). The resulting RF maps were thresholded by clipping negative values and fit with a 2D Gaussian. The retinotopic position of each neuron was estimated as the center of this Gaussian fit. To estimate preferred orientation, we presented full-contrast sinusoidal gratings across 36 orientations (0° to 175° in 5° steps), 36 phases (0° to 360°, in 10° steps), and 25 spatial frequencies (from 1.1 to 8.0 cycles per degree). For each neuron, we selected the orientation corresponding to the combination of phase and spatial frequency that elicited the highest response and defined that as the preferred orientation of the neuron.

#### Parameters and training of the machine learning models on macaque data

The macaque V1 population of neurons was divided into training and held-out sets according to the macaque of origin, with 238 neurons from the first macaque used for training and the remaining 214 neurons from the second macaque reserved for evaluating generalization performance. Input images were cropped to the center 2.67 degrees of visual angle (following the approach of Fu et al. [28] and **?**]), as all recorded neurons had receptive fields located in the central visual field, and subsequently downsampled by half to 46×46 pixels. To train the models, both stimuli and responses were z-scored: pixel values were standardized across the dataset to have zero mean and unit variance, and spike counts were z-scored independently for each neuron. The BRCNN core architecture consisted of 7 layers with input kernels of size 5 and hidden kernels of size 9. Each rotationally equivariant convolutional layer featured 8 channels (up to the final layer) and 32 orientations, followed by rotationally equivariant batch normalization and adaptive ELU nonlinearity. We used AdamW optimizer with an initial learning rate of 0.001 and batch size of 8, using early stopping when correlation stopped improving after 4 learning rate reductions. Hyperparameters were selected to maximize correlation between predictions and trial-averaged responses. The energy model was trained following the same approach, except that after z-scoring, input image pixel values were additionally shifted to be centered around 0.

#### Synthetic Spiking V1 Neural Network Dataset

The dataset used in the in-silico spiking network simulation experiments was collected by presenting grayscale images from ImageNet [21] to a previously validated large-scale spiking neural network model of V1 [3], described in earlier in the Methods section. It contains single-trial responses of the entire Layer 2/3 excitatory and inhibitory population (composed of 37,500 and 9,375 neurons, respectively) to 100,000 grayscale images, along with a smaller multi-trial subset consisting of 250 images used as the test dataset. Before being presented to the spiking neural network model, pixel values were rescaled to maximize the dynamic range of each image. To parallelize the simulations the dataset was divided into batches of 100 images, with each batch corresponding to a separate simulation. Each simulation began with an initialization period of 270 ms with no stimulation, followed by the sequential presentation of the 100 images. Each image was shown for 560 ms and followed by a 150 ms period of no stimulation before the next image, to avoid cross-contamination of responses. Individual neuron responses were collected by counting the number of spikes evoked during the presentation window for each image. The multi-trial part of the dataset was collected using a similar procedure. The 250 test images were divided across multiple simulations, each containing a subset of images presented for 100 trials in randomized order. As with the single-trial simulations, each began with an initialization period with no stimulation, and each image presentation lasted 560 ms, followed by 150 ms of no stimulation before the next image.

#### Spiking V1 neural network model neurons retinopic positions and preferred orientations

Neuron positions and preferred orientations used to fix the position and orientation parameters of the readout were obtained directly from the Spiking V1 Neural Network model, specifically, from the neurons position and orientation parameters used to define their connectivity rules, as described in Methods.

#### Parameters and training of the machine learning models on the synthetic V1 dataset

The neuron population was split into two groups: a training set (composed of 20000 excitatory and 5000 inhibitory neurons) and a held-out set (composed by the remaining neurons in the Layer 2/3 populations), used to evaluate the generalization performance of the BRCNN model’s core. Both stimuli and responses underwent z-scoring following the same procedure as the macaque V1 dataset.

Input images were downscaled by half to 55×55 pixels to improve computational efficiency. The BRCNN core was implemented with 5 layers, using input kernels of size 4 and hidden kernels of size 9. Each rotationally equivariant convolutional layer featured 4 channels (up to the final layer) and 32 orientations. Each layer was followed by a rotationally equivariant batch normalization operation and an adaptive ELU nonlinearity. Only the core component was trained during the optimization process (position and orientation parameters of neurons were kept fixed). The model was trained to minimize the mean squared error (MSE) between predictions and target neural responses. We used AdamW optimizer with an initial learning rate of 0.005, incorporating early stopping when ceased improving after two learning rate reductions. The batch size was set to 32 throughout training. Hyperparameters were choosing the configuration that maximized test dataset correlation between prediction and trial averaged responses on the held-out neuron set. The energy model was trained following the same approach, except that after z-scoring, input image pixel values were additionally shifted to be centered around 0.

### Maximally Exciting Images (MEIs)

MEIs [62] were obtained by optimizing input pixel values to maximize each neuron’s response, subject to constraints enforcing a mean equal to the mid-scale gray value and a fixed pixel-value standard deviation. This approach prevents the generation of features outside the neuron’s receptive field. The optimization consisted of performing gradient ascent on the same image for 1,000 steps with a learning rate of 10. To minimize artifacts, gradients were blurred using a Gaussian filter with *σ* = 1 pixels. After each optimization step, pixel values were linearly rescaled to satisfy the mean and standard deviation constraints. The image was then clipped to ensure that pixel values remained within the range corresponding to black and white in the training set. The pixel standard deviation was set to 0.15, chosen to minimize clipping while allowing the optimized images to exploit nearly the full dynamic range of pixel values.

### Size-tuning Experiments

Size-tuning experiments were performed by presenting full-contrast sinusoidal gratings within a circular mask of varying radius, centered at each neuron’s retinotopic position. Pixel values outside of the circular mask were set to mid-scale gray. Grating orientation and spatial frequency were matched to the neuron’s preferred values as determined in orientation tuning experiments. The stimulus radius ranged from 0.01° to 2° (in 9 logarithmically distributed steps) for macaque V1 neurons, and from 0° to 5.5° (in 40 steps) for spiking neural network neurons. Responses were averaged across all phases, and then across neurons, to generate the size-tuning curves shown in Fig.2D and Fig.3F.

### Stimulation Protocols

After training the BRCNN model, we used it to derive the stimulation protocol:

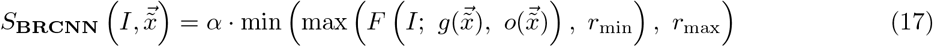

Specifically, given the high neuron density in the spiking V1 model, we computed the stimulation value for each electrode by identifying the neuron with the smallest lateral displacement relative to the electrode position, and using its predicted activation as 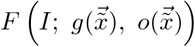. The output of the function *F* was clipped between *r*_min_ = 0 (the mean of the z-scored responses, setting the threshold for eliciting stimulation) and *r*_max_ = 5 (slightly above the model’s peak response under full-field grating stimulation and chosen to correspond to the maximum stimulation strength).

To calibrate *α*, we first presented four full-field oriented gratings (0°, 45°, 90°, and 135°) under normal vision conditions, each repeated 20 times, and recorded the corresponding neural responses. We then tested multiple values of *α* by applying optogenetic stimulation using the resulting patterns and measured the evoked responses. The final value of *α* was chosen as the one yielding the highest correlation between optogenetically evoked and visually evoked activity patterns. This calibrated value was subsequently used for both the BRCNN-based and the retinotopic stimulation protocols. This in-silico calibration procedure corresponds to calibrating the overall dynamical range of the stimulator in in-vivo experiments either based on neural readout or behavioral feedback.

The optogenetic experiments in Results were performed using these four oriented gratings and a set of 30 MNIST digits, similarly presented 20 times under both natural vision and optogenetic stimulation.

For each stimulus, we assessed stimulation fidelity by computing the Pearson correlation across neurons between the visually evoked and optogenetically evoked response vectors. This correlation served as a quantitative measure of how well the natural population response pattern was reproduced by each protocol.

## Acknowledgments

LB would like to thank Rèmy Cagnol for his valuable assistance and feedback on the technical implementation of the spiking V1 model. LB and JA were supported by the EU Horizon 2020 Maria SklodowskaCurie grant agreement No 861423. JA was supported by the ERDF-Project Brain dynamics, No. CZ.02.01.01 \00 \22 008 \0004643 and by the grant no. 25-18031S of the Czech Science Foundation (GAČR).

## Supplementary Material

### Large BRCNN overfit training neurons if dataset size is small

During hyperparameter search on macaque V1 data, we observed that increasing model depth and hidden kernel size improved accuracy on held-out images from the training neurons of one animal, but reduced generalization to neurons from a different animal. In a representative high-capacity configuration (7 layers; hidden kernel size 9), the trial-averaged Pearson correlation reached *r* = 0.52 on held-out images for training neurons, while generalization to unseen neurons dropped to *r* = 0.26. This pattern suggests overfitting to idiosyncrasies of the training neurons and highlights the need to match model size to dataset size and diversity, selecting capacity via cross-neuron or cross-subject validation rather than within-neuron performance alone.

## Footnotes

1 Note that this does not imply ON insensitivity; the BRCNN remains responsive to bright stimuli via its nonlinear computations.

2 This refers to the total number of images in the macaque dataset. For neurons from different recording sessions, different subsets of images and corresponding responses are available.

3 Despite this, it is worth pointing out that optogenetic stimulation has already successfully been tested in human retina in a recent clinical trial [55].

